# The p.H222P lamin A/C mutation induces heart failure via impaired mitochondrial calcium uptake in human cardiac laminopathy

**DOI:** 10.1101/2024.08.21.609073

**Authors:** Magali Seguret, Charlène Jouve, Andrea Ruiz-Velasco, Lucille Deshayes, Zoheir Guesmia, Céline Pereira, Karim Wahbi, Jérémy Fauconnier, Gisèle Bonne, Antoine Muchir, Jean-Sébastien Hulot

## Abstract

**Background:** Mutations in the *LMNA* gene, which encodes lamin A/C, cause a variety of diseases known as laminopathies. Some mutations are particularly associated with the occurrence of dilated cardiomyopathy and heart failure, but the genotype-phenotype relationship and underlying mechanisms are unclear. Here, we used induced pluripotent stem cells (hiPSCs) from a patient carrying a *LMNA* point mutation (c.665A>C, p.His222Pro) to investigate the mechanisms leading to contractile dysfunction.

**Methods:** LMNA p.H222P mutant and a CRISPR/Cas9 corrected isogenic control hiPSCs clones were differentiated into cardiomyocytes (hiPSC-CMs). Immunofluorescence staining was performed on hiPSC-CMs to quantify their sarcomere organization (SarcOrgScore) using a Matlab code. Ring-shaped cardiac 3D organoids were generated to compare the contractile properties of the two clones. Calcium transients in mutant and corrected hiPSC-CMs were measured by live confocal imaging. Mitochondrial respiration parameters were measured by Seahorse.

**Results:** hiPSC-CMs were generated from the *LMNA* mutant and the corrected hiPSCs with no difference in the differentiation yield (proportion of troponin-positive cells: 95.0% for *LMNA* p.H222P *vs*. 95.1% for Ctrl-iso1, p=0.726). hiPSC-CMs displayed well-formed sarcomeres and their organization was similar between the two cell lines. However, cardiac 3D organoids generated with *LMNA* p.H222P hiPSC-CMs showed an impaired contractility compared to control organoids. Calcium transient recordings in *LMNA* p.H222P mutant cardiomyocytes showed a significantly higher calcium transient amplitude with a significantly slower calcium re-uptake. Transcriptomic analyses suggested a global mitochondrial dysfunction and in particular an impaired mitochondrial calcium uptake with a significantly decreased expression of the mitochondrial calcium uniporter (MCU). This decrease in MCU expression was confirmed by western blot and was accompanied by an increased MICU1:MCU, as well as an increased PDH Ser232 and PDH Ser300 phosphorylation, indicating a decreased mitochondrial calcium uptake in the *LMNA* mutant hiPSC-CMs. Measurement of mitochondrial respiration showed lower basal and maximal respiration in *LMNA* p.H222P hiPSC-CMs. Consistently, the ATP levels were significantly lower in *LMNA* p.H222P hiPSC-CMs as compared to isogenic controls.

**Conclusions:** *LMNA* p.H222P mutant hiPSC-CMs exhibit contractile dysfunction associated with mitochondrial dysfunction with impaired MCU complex activity, decreased mitochondrial calcium homeostasis and reduced mitochondrial energy production.

**NOVELTY AND SIGNIFICANCE:** *What is known?:*

- Mutations in *LMNA*, which encodes the nuclear lamins A/C, cause a variety of diseases (called laminopathies), which can involve the cardiac muscle leading to dilated cardiomyopathy and systolic heart failure.
- The pathological mechanisms linking the nuclear envelope abnormalities induced by *LMNA* mutations to the development of a reduced cardiac muscle contractility are not well understood.

*What new information does this article contribute?:*

- *LMNA* mutant cardiomyocytes have a profound mitochondrial dysfunction with impaired MCU complex activity, decreased mitochondrial calcium homeostasis, and reduced mitochondrial energy production.
- Our study uncovers an unappreciated pathophysiological mechanism and opens new possibilities by suggesting MCU activators as a novel therapeutic for patients with *LMNA* cardiomyopathy.

## INTRODUCTION

Mutations in *LMNA*, which encodes the nuclear lamins A/C, cause a variety of diseases, called laminopathies. The majority of *LMNA* mutations are associated with muscular disorders^1^, which can include the cardiac muscle, resulting in *LMNA*-cardiomyopathy. Lamin A/C-related cardiomyopathy is one of the most common causes of familial dilated cardiomyopathy (DCM) and is characterized by a progression to heart failure in the early adulthood, associated with life-threatening cardiac arrhythmias, conduction defects and a high risk of sudden death^2, 3^. It is estimated that mutations in *LMNA* account for 5 to 8% of familial DCM, and DCM patients carrying *LMNA* mutations are now recognized as high-risk patients requiring specific guideline-recommended management including implantable cardioverter-defibrillator (ICD) implantation and early access to heart transplantation^3, 4^. However, the pathological mechanisms underlying the *LMNA*-associated cardiac phenotype are not well understood, thereby limiting the development of targeted therapeutic approaches.

Lamins are the major structural components of the nuclear lamina, a mesh-like structure located at the inner nuclear membrane^5^. Lamins support the mechanical integrity of the nucleus, are involved in chromatin organization, and interact with several other nuclear membrane proteins to modulate the cellular mechano-transduction and activation of signaling pathways^5^. Thus, *LMNA* mutations affect the nuclear envelope and have been shown to affect gene expression by inducing chromatin remodeling, or the mechanical response to stress by affecting the LINC (linker of nucleoskeleton and cytoskeleton) complex at the nuclear surface. For example, the *LMNA* p.K219T mutation has been shown to induce an epigenetic inhibition of *SCN5A*, the gene encoding the sodium channel, resulting in decreased sodium current density, slower conduction velocity, and conduction abnormalities^6^. In addition, *LMNA* mutations can alter the organization of actin and microtubule networks and the modulation of *α*-tubulin acetylation levels improved the cardiac function of *Lmna^H222P/H222P^*mice^7^. Additional functions are likely to be affected by *LMNA* mutations, supporting the transition to heart failure. An imbalance between oxidation and antioxidant response has been shown in *Lmna^H222P/H222P^*mice, with a decrease in glutathione peroxidase levels that was reversed by N-acetylcysteine administration^8^. In addition, a decrease in NAD^+^ levels was also found in *Lmna^H222P/H222P^* mice, and reversed by supplementing animals with the NAD^+^ precursor nicotinamide riboside (NR)^9^. Metabolic abnormalities may contribute to the pathogenesis of heart failure, but whether they are also important contributors to *LMNA*-related cardiomyopathy and heart failure is not firmly established.

In this study, we focused on the *LMNA* c.665A>C p.His222Pro mutation causing the Emery-Dreifuss Muscular Dystrophy (EDMD) which is characterized by joint contractures and muscle atrophy and weakness associated with dilated cardiomyopathy^10^. We aimed to develop a human cardiac model of the *LMNA* related cardiomyopathy using mutant and CRISPR/Cas9 corrected human induced pluripotent stem cells to elucidate the pathological abnormalities leading to the development of contractile dysfunction.

## MATERIAL AND METHODS

Details on the procedures are provided in the online-only data supplement. The data and material that support the findings of this study are available from the corresponding author upon reasonable request.

### Induced pluripotent stem cell lines

The hiPSC line heterozygous for the NM_170707.4(*LMNA*):c.665A>C (p.His222Pro) mutation was derived by Sendai virus infection from PBMCs of a 23-year-old male patient^10^ and is referred to as the *LMNA* p.H222P line. The non-isogenic SKIPSC-31.3 hiPSC line, derived from human dermal fibroblasts from a healthy 45-year-old control subject, as previously reported^11, 12^, was used as a reference for the functional assessments of the cardiomyocytes.

### Cardiomyocytes differentiation and culture

Mutant and corrected hiPSCs were differentiated when they reached 90% confluence, according to a protocol adapted from *Garg et al**.^13^***and previously described in other recent studies^12^, which includes a small-molecule driven mesorderm induction (using IWR-1 (Sigma)), and two rounds of glucose starvation to increase the percentage of cardiomyocytes obtained. After day 18, the cells were cultured in RPMI-B27 with insulin and the medium was changed every two days.

### 3D cardiac rings generation and recording

3D cardiac rings were generated as previously described^14^. Briefly, the tissues are generated on a 96-well plate containing a molded hydrogel in each well that induces seeded cells to self-organize into up to 21 ring-shaped tissues per well. At D22 of differentiation, cardiomyocytes from *LMNA* p.H222P mutant and corrected cell lines were dissociated and mixed with dissociated healthy human dermal fibroblasts at a hiPSC-CM:fibroblast ratio of 3:1, to produce mutant or corrected cardiac tissues, respectively. At 10 days after their generation, the tissues were recorded in brightfield imaging with a high-speed CCD camera (PLD672MU, Pixelink) mounted on a microscope (Primovert, Zeiss), at a 10X magnification. Automated video analysis was performed using a Matlab script, which allowed the recovery of contractility parameters such as the contraction frequency, the contraction stress, and the maximum contraction and relaxation speeds

### Statistical analysis

All numerical results are expressed as mean ± standard deviation (SD) of three independent experiments. The number of samples (*n*) used in each experiment is recorded in the text and figure legends. Quantitative variables were analyzed with the Mann-Whitney U test for comparison of data obtained in the two studied groups, *LMNA* p.H222P vs. the isogenic control (Ctrl-Iso1). Data from the unrelated healthy hiPSC line are reported as a reference but are not statistically compared to the other two groups. P values < 0.05 were considered significant for all statistical tests. Statistical analyses were performed with GraphPad Prism software.

## RESULTS

### Generation and characterization of *LMNA* p.H222P mutant and corrected hiPSCs-derived cardiomyocytes

We used a human induced pluripotent stem cell line generated from a patient heterozygous for the *LMNA* c.665A>C mutation. This missense variant leads to a change in the amino acid sequence (p.His222Pro) associated with an autosomal dominant Emery-Dreifuss Muscular Dystrophy (EDMD2) and LMNA-related cardiomyopathy as described previously^10^. To obtain an isogenic corrected clone, we used CRISPR/Cas9 to correct the *LMNA* c.665A>C mutation. A mutation-specific gRNA was designed. We found that the presence of the C variant on position 665 of the *LMNA* gene induces the generation of a PAM sequence that can be specifically recognized by the CRISPR/SpCas9 system (**Figure 1A**). We then designed an optimized donor matrix to correct the DNA through homologous directed repair, with silent mutations to the recombination matrix to prevent unwanted re-cutting of the corrected DNA as well as gRNA homology on the donor matrix, in line with previous reports^15^ (**Figure 1A** and supplementary table 1). We used the ESEfinder software^16, 17^ to ensure that the synonymous SNPs would not induce any major modification of the exonic splicing enhancers in the vicinity of the mutation. The gRNA sequence was inserted into a plasmid containing the sequence of SpCas9 and nucleofected with the donor matrix. The electroporated clones were isolated and sequenced, and a corrected clone showing correction of the *LMNA* c.665A>C mutation was selected (**Figure 1B**), and referred to as Ctrl-iso 1. We excluded the presence of off-target events by sequencing the most likely predicted off-target sites, in the corrected and mutant clones (Supplemental Figure 1). The pluripotency of Ctrl-iso 1 hiPSCs was verified by immunofluorescence (Supplemental Figure 2A., and quantitative PCR of pluripotency genes (*Nanog*, *Sox2*, *Oct4* and *TERT*) (Supplemental Figure 2B). No karyotype abnormality was observed in both hiPSC clones (Supplemental Figure 2C). We also verified that the correction plasmid sequence was not integrated into the DNA of the corrected clone (Supplemental Figure 2D).

**Figure 1:**
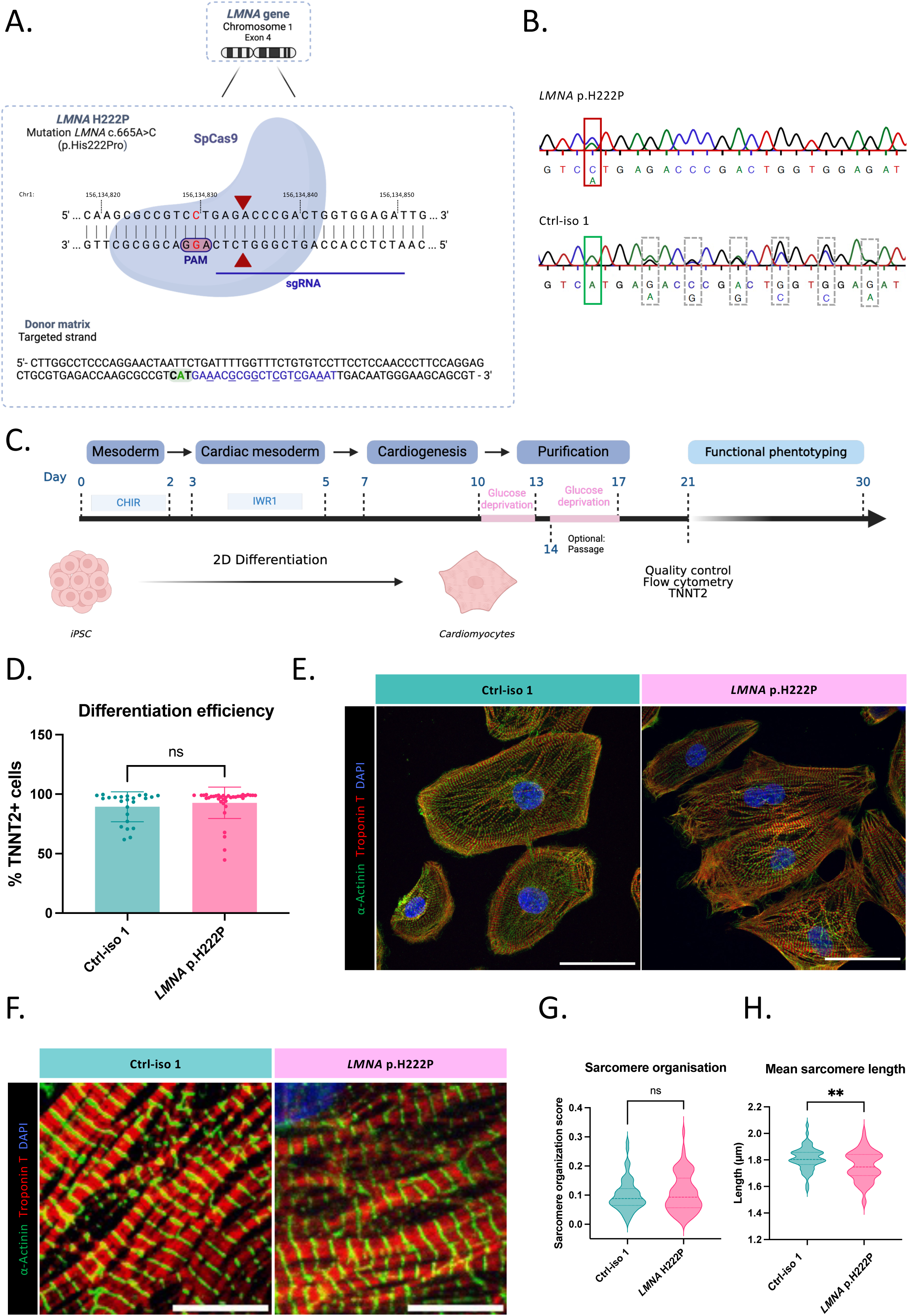
Correction of the *LMNA* c.665A>C (p.His222Pro) mutation in heterozygous hiPSCs and differentiation in cardiomyocytes. **A.** CRISPR-Cas9 strategy used for the correction of the *LMNA:* c.665A>C (p.His222Pro) mutation (nucleotide in red). The gRNA sequence used to cut the DNA in the vicinity of the mutation is underlined in blue on the 3’ strand. The PAM sequence (AGG) on the 3’ strand is mutation-specific. The donor matrix was designed on the targeted strand (36 nucleotides on the PAM-distal side, 91 nucleotides on the PAM-proximal side) and displays the corrected sequence (corrected nucleotide in green) and 6 synonymous SNPs (underlined). The part of the donor matrix sequence which Is complementary to the gRNA is in blue. **B.** Sequences of the *LMNA* p.H222P mutant hiPSC clone and of the corrected clone Ctrl-iso. The heterozygous c665A>C mutation (red frame in the mutant clone) is not found in the corrected clone (green frame), which is heterozygous for the synonymous SNPs inserted in the donor matrix, (ramed in grey). **C.** Protocol of differentiation used to obtain cardiomyocytes from hiPSCs. **D.** Differentiation efficiency expressed as the percentage of troponin T-positive cells measured by flow cytometry 21 days after the beginning of differentiation in the *LMNA* p.H222P mutant. Vs Ctrl-Iso1. Mean ± SD, N=26 and 37 differentiations respectively for Ctrl-iso 1 and *LMNA* p.H222P. Mann-Whitney test between *LMNA* p.H222P and Ctrl-iso 1. **E.** Representative staining of alpha-actinin (green), troponin T (red) and DAPI (blue) in Ctrl-iso 1 and *LMNA* p.H222P hiPSC-CMs, 30 days after the beginning of differentiation. Scale bar 50 µm. **F.** Pictures of sarcomeres immunostaining (alpha-actinin in green, troponin T in red) of Ctrl-iso 1 and *LMNA* p.H222P hiPSC-CMs. Scale bar 10 µm. **G-H.** Sarcomere organization score (G) and sarcomere length (H) determined with a Matlab code for Ctrl-iso 1 and *LMNA* p.H222P. n= 57 and 68 cells respectively for Ctrl-iso 1 and *LMNA* p.H222P, for 2 differentiations. Mann-Whitney test.

Both *LMNA* p.H222P mutant and isogenic control cell lines can be differentiated into cardiomyocytes with a similar efficiency, as shown by quantification of TNNT2-positive cells by flow cytometry (**Figure 1C-D**). Immunostaining for alpha-actinin and troponin T at day 30 of differentiation shows well-formed sarcomeres in all cell lines, which are characteristics of cardiomyocytes (**Figure 1E**). To further analyze the contractile apparatus of the two cell lines, we used a Matlab script to quantify the sarcomere organization and the sarcomere lengths. We found no difference in sarcomere organization score the *LMNA* mutant and corrected cell lines (**Figures 1F-G**). The sarcomere length was however lower in *LMNA* p.H222P hiPSC-CMs as compared to corrected cells, with a trend for a bimodal distribution in the *LMNA* p.H222P hiPSC-CMs. Therefore, differentiation of mutant and corrected hiPSC lines results in apparently well-formed cardiomyocytes.

### *LMNA*-mutant hiPSC-derived cardiomyocytes show nuclear abnormalities

A Matlab script was used to quantify additional morphological features of hiPSC-derived cardiomyocytes. We also used an unrelated hiPSC derived from a healthy donor (referred to as WT) as another reference. No difference in the cell area was observed between *LMNA* p.H222P and Ctrl-iso 1 cell lines (**Figures 2A-B**). These parameters were comparable to those of the non-isogenic WT hiPSC-CMs. Since the *LMNA* mutation affects the nuclear membrane, we then compared different nuclear morphology descriptors. We found that the nuclei of *LMNA* p.H222P mutant hiPSC-CMs were significantly larger but less elongated compared to isogenic corrected hiPSC-CMs in 2D cell monolayers (**Figures 2A, C-D**). Importantly, lamin A/C expression was similar in cardiomyocytes derived from mutant and corrected cell lines (**Figure 2E-G**), and similar to the non-isogenic WT hiPSC-CMs.

**Figure 2:**
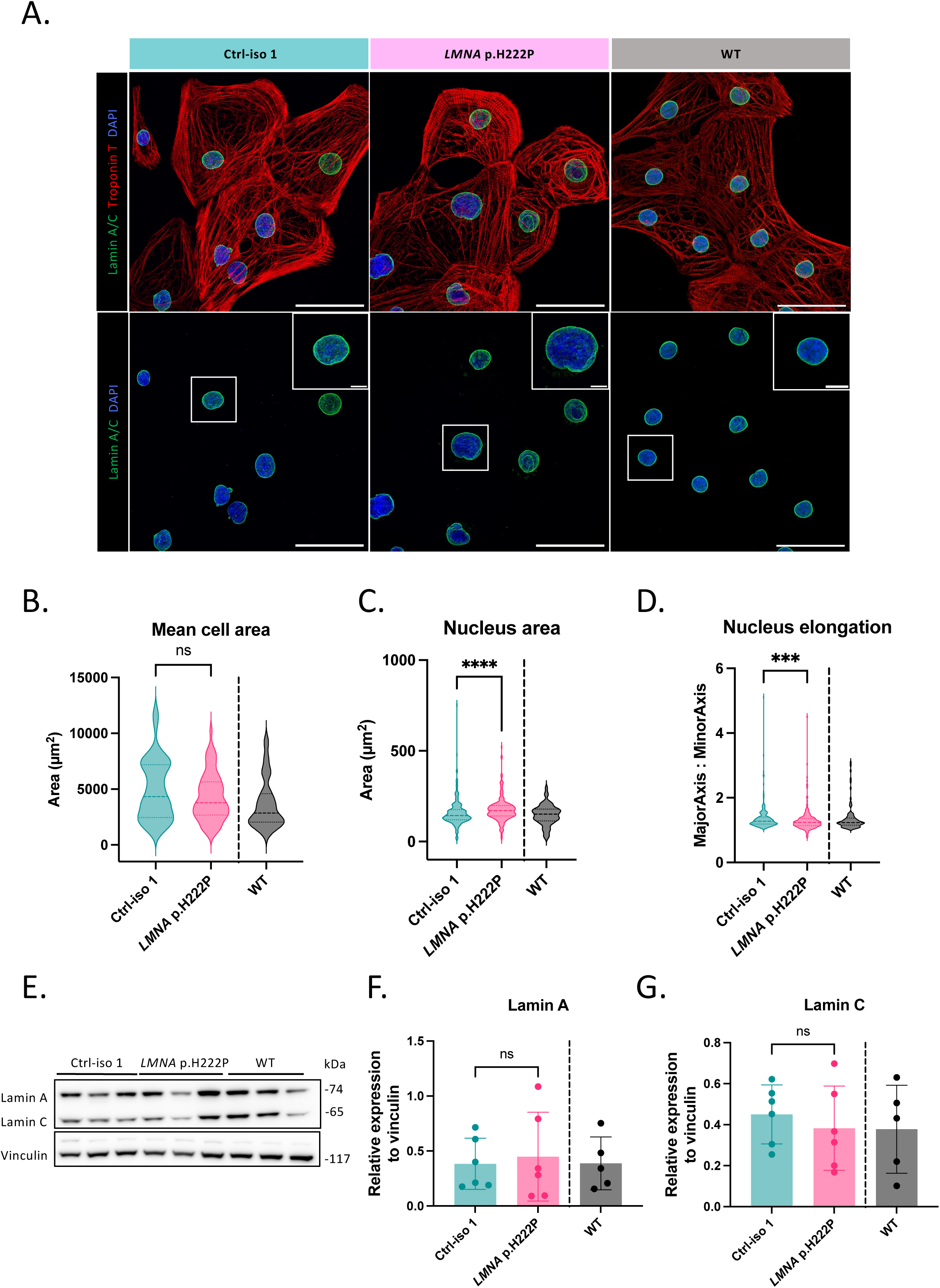
Cardiomyocytes morphological characterization. **A.** Immunostaining of lamin A/C (green), troponin T (red) and DAPI (blue) in *LMNA* p.H222P and Ctrl-iso 1 hiPSC-CMs 30 days after the beginning of differentiation. Merge is shown in the upper panel and lower panel shows immunostaining of lamin A/C (green) and DAPI (blue). The insert represents a zoomed view of a nucleus. Global scale bar 50 µm, insert scale bar 10 µm. **B.** Quantification of mean cell area in µm^2^ measured on immunostainings for the Ctrl-iso 1, *LMNA* p.H222P and non-isogenic WT hiPSC-CMs. n= 57, 68 and 20 cells respectively for Ctrl-iso 1, *LMNA* p.H222P and non-isogenic WT hiPSC-CMs for 2 differentiations. Mann-Whitney test between *LMNA* p.H222P and Ctrl-iso 1. **C.** Nucleus area and **D.** elongation quantifications in mutant and corrected hiPSC-CMs, calculated with a bioinformatic routine. n= 223, 227 and 98 nuclei respectively for Ctrl-iso 1, *LMNA* p.H222P and non-isogenic WT hiPSC-CMs. Mann-Whitney test between Ctrl iso 1 and *LMNA* p.H222P. **E.** Western blot of lamin A/C and vinculin on Ctrl-iso 1, *LMNA* p.H222P and non-isogenic WT hiPSC-CMs protein extracts. N=3 differentiations for each cell line. **F.** Quantification of lamin A and lamin C **G.** expression in Ctrl-iso 1, *LMNA* p.H222P and non-isogenic WT hiPSC-CMs, expressed as relative to vinculin expression for each sample. Mean ± SD, N=3 differentiations for each cell line. Mann-Whitney test between *LMNA* p.H222P and Ctrl-iso 1.

### Engineered cardiac tissues generated with *LMNA* p.H222P mutant hiPSC-CMs display contractile dysfunction

We then evaluated the functional characteristics of *LMNA* mutant and corrected hiPSC-CMs. Since the function is influenced by the 3D environment, including the extracellular matrix and the multicellular interactions, we generated engineered cardiac tissues (hECT) using the *LMNA* mutant, corrected and unrelated healthy hiPSC-CMs respectively. Briefly, hiPSC-CMs were mixed with human dermal fibroblasts at day 22 of CM differentiation and seeded in an assay to generate self-assembled 3D cardiac rings (**Figure 3A**), as previously described^14^. The rings were imaged 10 days later with a high-speed camera and the contractile properties of the cardiac tissues were assessed by tracking the area of the central pillar over time on the brightfield videos of the rings. We first observed that both *LMNA* p.H222P and Ctrl-iso 1 engineered cardiac tissues were well formed and had a similar morphology and showed contractile activity (**Figure 3B-C, and supplemental movies 1-2**). However, the engineered cardiac tissues generated with the *LMNA* p.H222P mutant hiPSC-CMs exhibited an impaired contractility with significantly lower contraction frequency, lower contraction stress and slower maximum contraction and relaxation velocities (**Figure 3C**), as compared to the cardiac tissues generated with the corrected or the WT hiPSC-CMs. Conversely, the *LMNA* corrected hiPSC-CMs showed a restored contractility, similar to that measured in the non-isogenic WT control (**Figure 3C**). Consistent with our results showing similar sarcomeric organization between the cell lines (**Figure 1E-F**), immunofluorescence staining for troponin T showed that the tissues had a similar and well-formed internal structure (**Figure 3D**). In addition, we analyzed the nuclear morphology of hiPSC-CMs in the engineered cardiac tissues and found significant differences with smaller and more elongated nuclei in tissues generated with the *LMNA* p.H222P mutant hiPSC-CMs (**Figure 3E**). Taken together, these results indicate that the *LMNA* p.H222P mutation affects the structure of the nuclear envelope, but does not affect the ability to generate cardiomyocytes in a 2D monolayer. However, the *LMNA* p.H222P mutation is associated with a significant reduction in contractility and relaxation as demonstrated in engineered cardiac tissues, and induce a significant effect on nuclear morphology in such a 3D environment.

**Figure 3:**
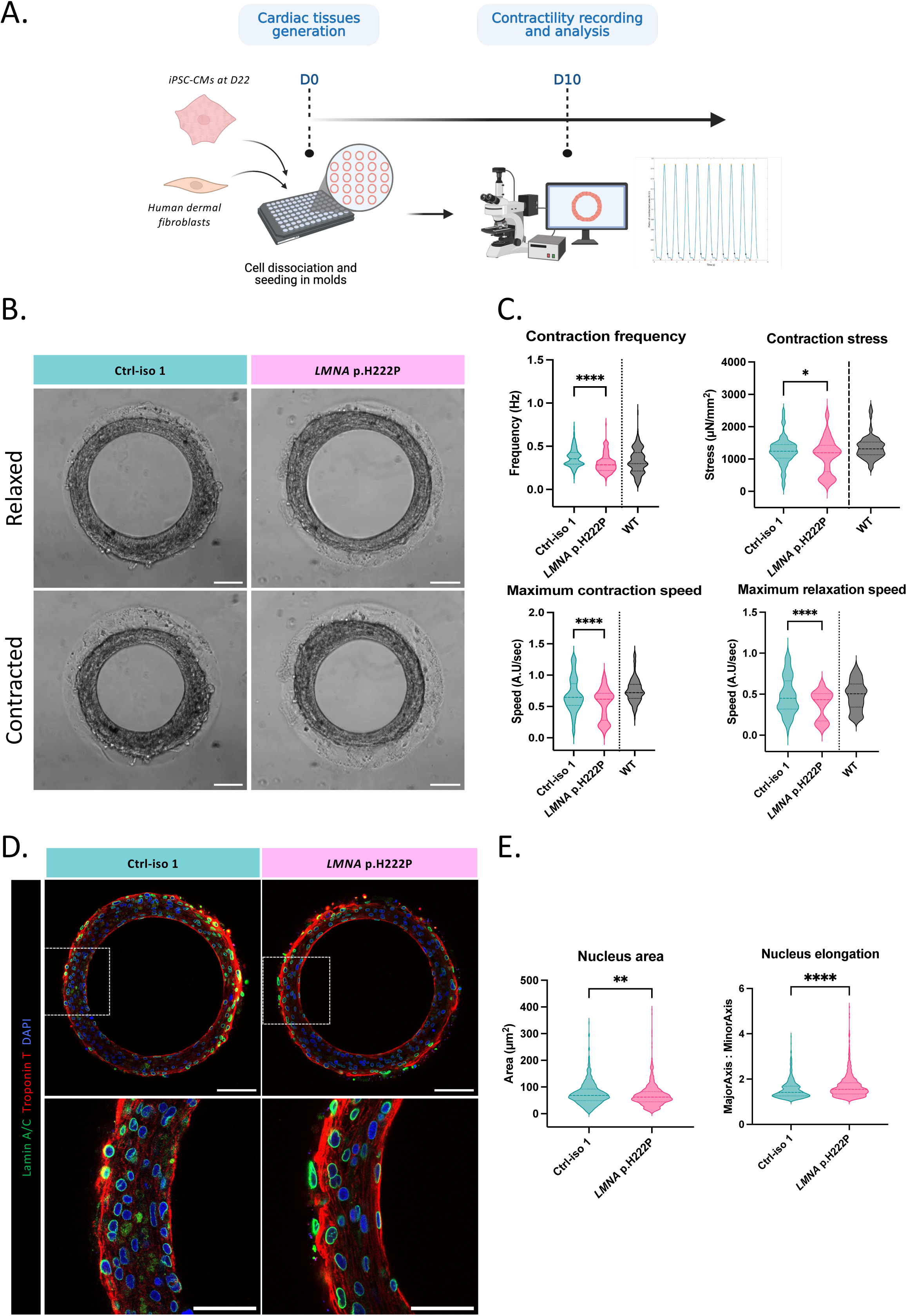
Tissue contractility. **A.** Timeline for the tissue generation and the video recordings. **B.** Brightfield images of representative tissues generated with *LMNA* p.H222P or Ctrl-iso 1 cardiomyocytes, in their most relaxed (upper picture) or contracted (lower picture) state. Scale bar 100 µm. **C.** Contractile parameters of *LMNA* p.H222P and Ctrl-iso1 tissues, compared to a non-isogenic control. n=164, 152 and 183 tissues respectively for Ctrl-iso 1, *LMNA* p.H222P and non-isogenic WT. Mann-Whitney test between *LMNA* p.H222P and Ctrl-iso 1. ****: p<0.0001; *: p<0.05. **D.** Troponin (red), lamin A/C (green) and DAPI (blue) staining of Ctrl-iso 1 and *LMNA* p.H222P cardiac tissues. Scale bars 100 µm. **E.** Nucleus area and elongation quantification in mutant and corrected 3D tissues. n=589 and 833 nuclei respectively for Ctrl-iso 1 and *LMNA* p.H222P 3D cardiac tissues. Mann-Whitney test between Ctrl-iso 1 and *LMNA* p.H222P.

### *LMNA* p.H222P hiPSC-CMs have abnormal calcium transients, reversed by the correction of the mutation

Cardiomyocyte contractility is closely related to intracellular calcium activity. Therefore, we evaluated calcium transients in our *LMNA* mutant and corrected hiPSC-CMs at 30 days of differentiation by live imaging with the fluorescent dye Fluo4-AM (**Figure 4A**). HiPSC-CMs were efficiently paced at 1Hz, with no significant difference between the different cell lines (**Figure 4B**). To remove one source of variability, only cells with a contraction frequency between 0.9Hz and 1.1Hz were analyzed. Therefore, we compared cells in the same range of beating frequency and compared their calcium traces (**Figure 4C**). Strikingly, we observed a significantly increased calcium transient amplitude in *LMNA* mutant hiPSC-CMs, with a marked increase in the area under the ΔF/F_0_ curve for *LMNA* p.H222P mutant cardiomyocytes, a signal that was reversed in Ctrl-iso 1. In addition, *LMNA* p.H222P cardiomyocytes exhibit a significant decrease in maximum calcium decay speed, whereas the rate of rise of the calcium transient was not affected by the mutation (**Figure 4D**). The expression of several genes involved in calcium transients (*CACNA1C, RYR2, SERCA2A, PLN, CASQ2*) was measured and no significant difference was found between *LMNA* p.H222P *vs*. Ctrl-iso (**Figure 4E**). These results strongly suggest that the defective contractility observed in the *LMNA* p.H222P mutant hiPSC-CMs involves an abnormal calcium cycling, which is not explained by an abnormal calcium handling by the sarcomeric reticulum (SR).

**Figure 4:**
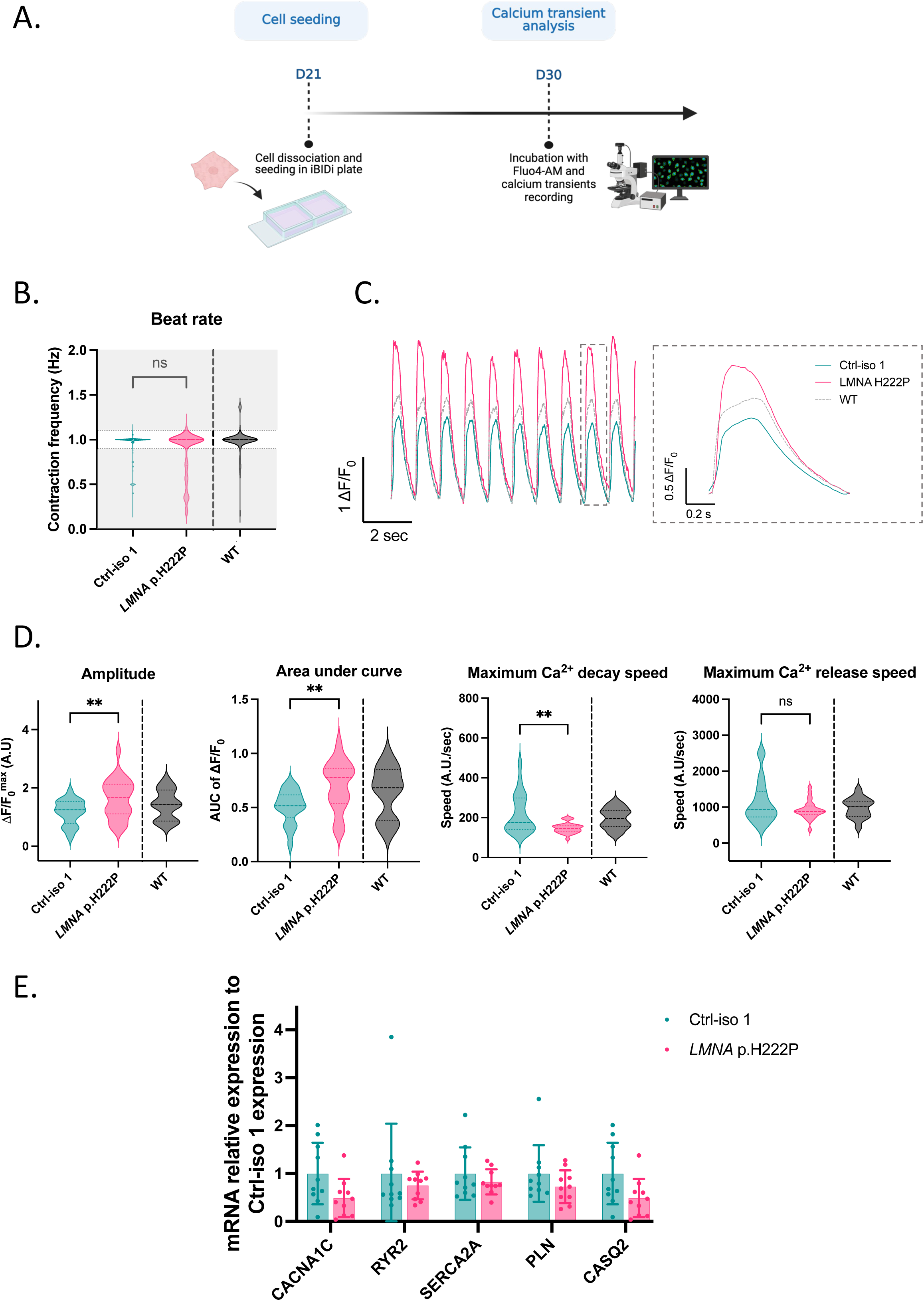
Calcium transient analysis. **A.** Timeline for calcium transient measurements. **B.** Calcium transient frequency (in Hz) of Ctrl-iso 1, *LMNA* p.H222P and non-isogenic WT hiPSC-CMs. n=203, 232 and 174 cells respectively for Ctrl-iso 1, *LMNA* p.H222P and WT non-isogenic control hiPSC-CMs. Only the hiPSC-CMs within the 0.9-1.1 Hz range (dotted lines) were selected for calcium transient analysis. **C.** Left: Representative traces of Ctrl-iso 1, *LMNA* p.H222P and non-isogenic WT hiPSC-CMs. Scale: 1ΔF/F0 for Y-axis, 2 seconds for X-axis. Right: Representative calcium transient trace for each cell line. Scale: 0.5ΔF/F0 for Y-axis, 0.2 seconds for X-axis. **D.** Calcium transient parameters of hiPSC-CMs mutant or corrected for the *LMNA* p.H222P mutation, and incubated with Fluo4-AM. Data points consist in the averaged parameter for a field: n=21, 26 and 16 fields respectively for Ctrl-iso 1, *LMNA* p.H222P and non-isogenic WT, for 3 independent experiments. Mann-Whitney test between *LMNA* p.H222P and Ctrl-iso 1 hiPSC-CMs. **: p<0.005; ***: p<0.001. **E.** mRNA expression of genes related to the SR calcium handling in Ctrl-iso 1 and *LMNA* p.H222P hiPSC-CMs: *RYR2*, *SERCA2A*, *CACNA1C*, *CASQ2* and *PLN* gene expression in hiPSC-CMs as a relative to *RPL32* expression. Mean ± SD, N= 10 differentiations for each clone. Mann-Whitney test between *LMNA* p.H222P and Ctrl-iso 1.

### Transcriptomic analysis highlights metabolic differences between *LMNA* p.H222P hiPSC-CMs and their isogenic control

In order to decipher the underlying mechanisms, transcriptomic analysis of the *LMNA* p.H222P mutant and Ctrl-iso 1 hiPSC-CMs was performed. By comparing gene expression in *LMNA* p.H222P mutant hiPSC-CMs to Ctrl-iso1 hiPSC-CMs, we found significant differences with 24 up-regulated and 14 down-regulated genes (**Figure 5A**). To better correlate these expression changes with the function of hiPSC-CMs, pathway analysis was performed using ReactomeGSA. Among the differentially expressed genes, many of them were zinc finger proteins, proteins involved in cell cycle and transcription/translation, up-regulation of JNK and Notch signaling pathways, in line with previous reports in the literature for laminopathic cardiomyocytes^18, 19^ (Supplemental Figure 3). However, among 43 significantly altered pathways, we found that 20 of these pathways were related to genes involved in metabolism (15 pathways), or protein metabolism (5 pathways) (**Figure 5B**). **Figure 5C** shows the significantly altered (p*_value_*<0.05) metabolic pathways in *LMNA* p.H222P mutant hiPSC-CMs compared to their isogenic control. In particular, we observed an impairment in the response to oxidative stress, including an upregulation of the beta-oxidation of very long chain fatty acids, lipophagy and biological oxidations. On the other hand, the pathways associated with processes leading to ATP production, such as intracellular fatty acid transport and the electron transport chain, were downregulated (**Figure 5D**). Several genes associated with the mitochondrial function were dysregulated in *LMNA* p.H222P hiPSC-CMs. We notably found a significant reduction in the expression of mitochondrial calcium uniporter (*MCU*) a transmembrane protein that allows the passage of calcium ions from the cytosol into the mitochondria, in the *LMNA* p.H222P mutant cells (**Figure 5E**). Similarly, we found a significant reduction in genes associated with mitophagy (*SPATA18*), electron transport chain activity (*FDXR*) and a stabilizing component of the Complex I of the mitochondrial respiratory chain (*NDUFAF7*) (**Figure 5E**). Conversely, we found no change in the expression of key cardiac genes (Supplemental Figure 4). We then further interrogated a transcriptomics profiling previously performed in murine hearts from *Lmna*^H222P/H222P^ transgenic mice vs. controls^7^. Consistent with our results from hiPSC-CMs, GO analysis showed that metabolic pathways were significantly dysregulated in hearts from *Lmna*^H222P/H222P^ transgenic mice, including fatty acid metabolism and TCA cycle (Supplemental Figure 5A). In line with our results focusing on genes involved in mitochondria, we also observed a significant reduction in the expression of *Mcu* and *Ndufaf7* (Supplemental Figure 5B). Overall, these results indicate a significant alteration in genes involved in mitochondrial process.

**Figure 5:**
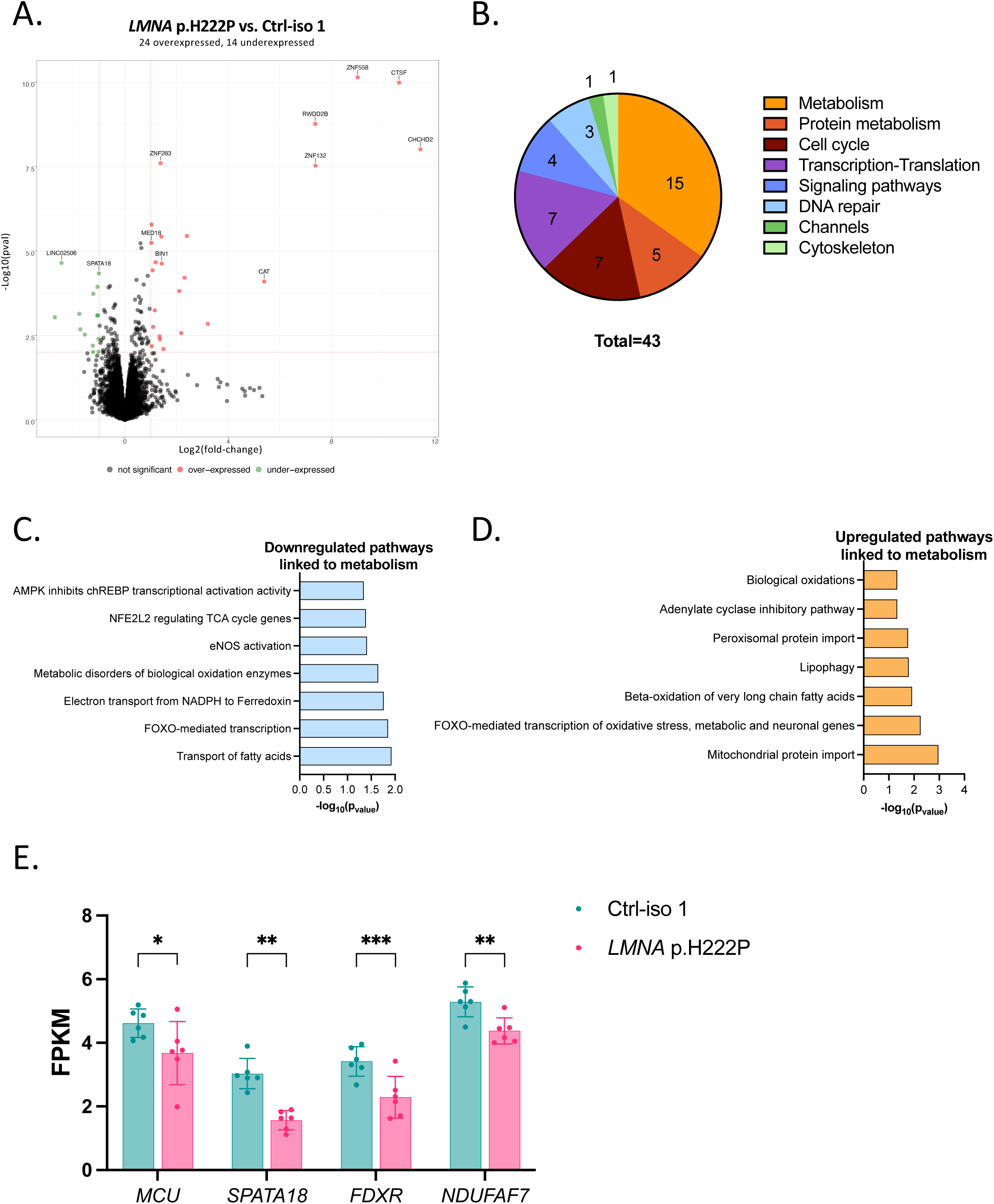
Transcriptomics and pathway analyses in *LMNA* p.H222P mutant hiPSC-CMs in comparison to their isogenic corrected control. **A.** Volcano plot of down-regulated (14, in green) and up-regulated (24, in red) genes in the *LMNA* p.H222P hiPSC-CMs in comparison to their corrected isogenic control Ctrl-iso 1 hiPSC-CMs. FPKM >1; Fold change >2 and p value<0.01. **B.** Classification of significantly changed pathways (p value<0.05) in *LMNA* p.H222P vs. Ctrl-iso 1 hiPSC-CMs with ReactomeGSA pathway analysis. **C-D.** significantly downregulated (C) and up-regulated (D) pathways related to energy metabolism (pvalue<0.05) in *LMNA* p.H222P vs. Ctrl-iso 1 hiPSC-CMs, as determined with ReactomeGSA pathway analysis. **E.** Differential gene expression of genes related to the quality and efficiency of mitochondria: *MCU*, *SPATA18*, *FDXR* and *NDUFAF7*. *: p<0.05; **: p<0.005; ***: p<0.001. Mean ± SD, N=6 differentiations.

### *LMNA* p.H222P mutant hiPSC-CMs show an impaired mitochondrial calcium uptake

Following the transcriptomic analyses suggesting mitochondrial dysfunction in *LMNA* mutant iPSC-CMs, we further examined the expression of proteins composing the mitochondrial calcium uniporter complex (MCUC) (**Figure 6A**). Consistent with the RNA sequencing results, western blot analyses confirmed the significant reduction in MCU expression in *LMNA* p.H222P mutant hiPSC-CMs (**Figure 6B-C**). In addition, we found a significant increase in the expression of the regulatory subunit MICU1 (**Figure 6B-D**), resulting in a strong increase in the MICU1:MCU (4.332 ± 3.178 *vs*. 1.521 ± 1.179, p<0.05, **Figure 6D**). MICU2 expression was however unchanged between the groups (Supplemental Figure 6).

**Figure 6:**
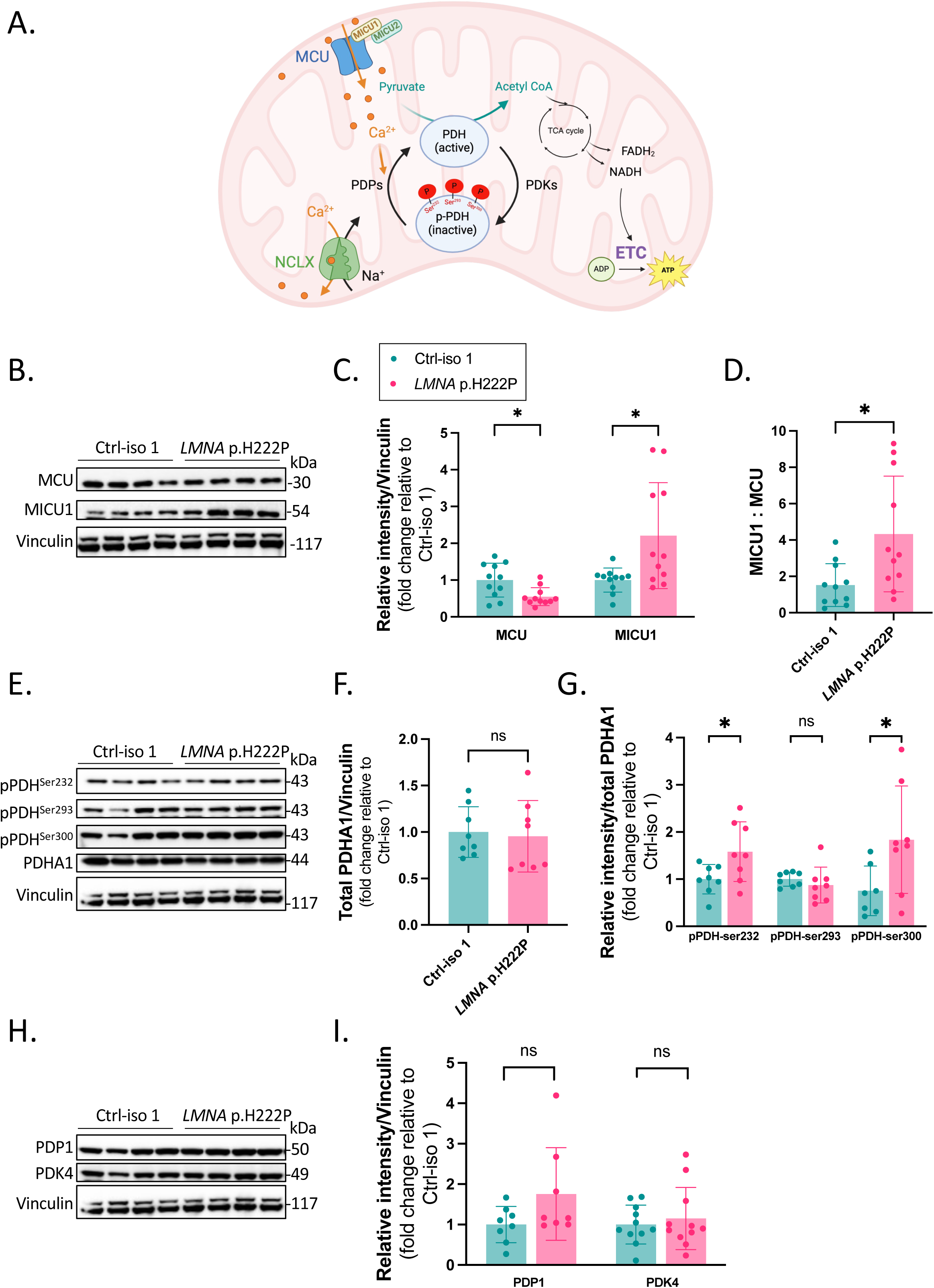
Impaired mitochondrial calcium handling in *LMNA* p.H222P mutant compared to Ctrl-iso 1 hiPSC-CMs. **A.** Scheme depicting the main actors of the mitochondrial calcium uptake and processes involved in energy production in mitochondria: MCU in blue, NCLX in green, active and inactive PDH with its phosphorylation sites in red. **B.** Representative immunoblots for MCU and MICU1. **C.** Quantification of relative protein level of MCU and MICU1 in hiPSC-CMs isolated from Ctrl-iso 1 and *LMNA* p.H222P. Vinculin was used as loading control and expressed relative to Ctrl-iso 1. Mean ± SD, N = 11 differentiations. Mann-Whitney test between *LMNA* p.H222P and Ctrl-iso 1. *: p<0.05 **D.** Proteins ratios of MICU1 to MCU, calculated individually for each differentiation. Unpaired t test between *LMNA* p.H222P and Ctrl-iso 1. *: p<0.05. **E.** Representative immunoblots for total pyruvate dehydrogenase subunit E1**α** (PDH-E1**α**), PDH phosphorylated on Serine 232, 293, and 300. **F.** Quantification of relative protein level of PDH-E1*α* in hiPSC-CMs isolated from Ctrl-iso 1 and *LMNA* p.H222P. Vinculin was used as loading control and expressed relative to Ctrl-iso 1. Mean ± SD, N=8 differentiations. Unpaired t test between *LMNA* p.H222P and Ctrl-iso 1. **G.** Quantification of relative protein level of PDH phosphorylated on Serine 232, 293 and 300 over total PDH-E1 *α* in hiPSC-CMs isolated from Ctrl-iso 1 and *LMNA* p.H222P and expressed relative to Ctrl-iso 1. Mean ± SD, N = 8 differentiations. Unpaired t test between *LMNA* p.H222P and Ctrl-iso 1. *: p<0.05. **H.** Representative immunoblots for PDP1 and PDK4. **I.** Quantification of relative protein level of PDP1 and PDK4 in hiPSC-CMs isolated from Ctrl-iso 1 and *LMNA* p.H222P. Vinculin was used as loading control and expressed relative to Ctrl-iso 1. Mean ± SD, N=8-11 differentiations. Mann-Whitney test between *LMNA* p.H222P and Ctrl-iso 1.

The increased MICU1:MCU has previously been associated with reduced mitochondrial calcium uptake^20^. To test this, we measured the pyruvate dehydrogenase (PDH) phosphorylation in *LMNA* p.H222P and Ctrl-iso1 hiPSC-CMs. PDH phosphorylation is modulated by the mitochondrial calcium concentration as dephosphorylation of PDH is catalyzed by the pyruvate dehydrogenase phosphatase 1 (PDP1), whose activity is enhanced when mitochondrial calcium levels increase (**Figure 6A**). We first verified that the total PDH content was unchanged (**Figure 6E-F**), as well as PDP1 and PDK4 expressions (**Figure 6H-I**). The three phosphorylable serines of PDH were then evaluated: Ser 232, Ser 293 and Ser 300. We found significant increases in the pPDHSer^232^:PDHtotal and pPDHSer^300^:PDHtotal ratios in *LMNA* p.H222P mutant hiPSC-CMs (**Figure 6G**). Moreover, the expression of the mitochondrial Na^+^/Ca^2+^ exchanger (NCLX) was unchanged (Supplemental Figure 6).

Therefore, together with the previous observations, these results highlight the remodeling of the MCUC in *LMNA* p.H222P mutant hiPSC-CMs, leading to decreased mitochondrial calcium uptake and calcium-dependent activation of mitochondria as a pathogenic feature that appears from the early stage of the *LMNA*-related cardiomyopathy.

### *LMNA* p.H222P hiPSC-CMs show an impaired mitochondrial respiration

As PDH activity modulates the production of Acetyl-CoA, which is implicated in the TCA cycle which produces substrates used in the electron transport chain (ETC) for ATP production, we next evaluated the electron transport chain activity in the *LMNA* p.H222P mutant hiPSC-CMs in comparison to their isogenic control. First, we measured the mitochondrial volume by flow cytometry and did not find any difference between the corrected and mutant hiPSC-CMs (**Figure 7A-B**). The assessment of mitochondrial respiration activity was then carried out with a Seahorse XFp analyzer. Thirty-five days after the beginning of differentiation, hiPSC-CMs were seeded on a micropatterned substrate, which was found to improve their metabolism^21^, and were transferred into Seahorse plates 7 days after, and analyzed with a Seahorse flux analyzer at D46 (**Figure 7C**). hiPSC-CMs were sequentially submitted to (1) the ATP synthase inhibitor oligomycin (2) the respiratory uncoupler FCCP, and (3) the respiratory chain blockers rotenone and antimycin A (**Figure 7D**). This allowed us to derive the basal and maximal respirations of the cell, as well as the ATP production. We found that basal and maximal respirations were significantly impaired in *LMNA* p.H222P cells, as well as the ATP-linked respiration (**Figure 7E**). The proton leak, the coupling efficiency, the spare respiratory capacity and the non-mitochondrial oxygen consumption were unchanged (Supplemental Figure 7). We finally measured the ATP content with a luminescence ATP detection assay system (**Figure 7 F**), and found significantly lower levels in *LMNA* mutant cells as compared to isogenic controls, thus confirming the energy production deficit in the LMNA mutant hiPSC-CMs.

**Figure 7:**
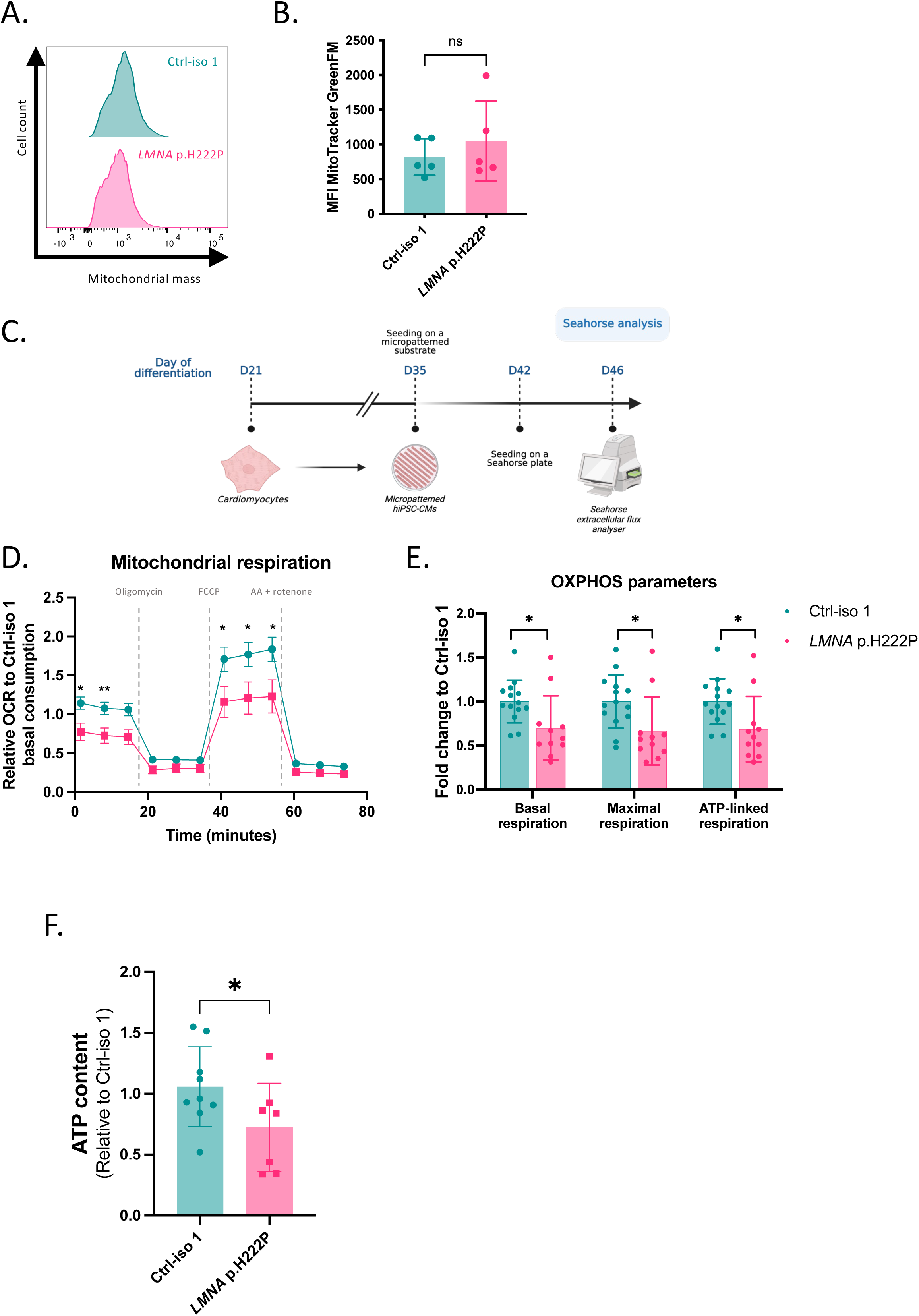
Impaired OXPHOS activity in *LMNA* p.H222P mutant compared to Ctrl-iso 1 hiPSC-CMs. **A.** Representative flow cytometry images from Ctrl-iso 1 and *LMNA* p.H222P hiPSC-CMs stained with MitoTracker™Green FM fluorescent dye. **B.** Quantification of the Mean Fluorescence Intensity of MitoTracker™Green FM from Ctrl-iso 1 and *LMNA* p.H222P hiPSC-CMs. Mean ± SD, N=5 differentiation with triplicates. ns p>0.05, Mann-Whitney test. **C.** Workflow of the mitochondrial respiration measurement on corrected (Ctrl-iso 1) and mutant (*LMNA* p.H222P) hiPSC-CMs. **D.** Real-time oxygen consumption rates measurements in *LMNA* p.H222P and Ctrl-iso 1 hiPSC-CMs by Seahorse extracellular flux analyzer. Cells were sequentially submitted to several drugs: (1) the ATP synthase inhibitor oligomycin (2) the respiratory uncoupler FCCP (3) the respiratory chain blockers rotenone and antimycin A. Mean ± SD, n=10 wells for *LMNA* p.H222P and n=13 wells for Ctrl-iso 1, from 3 independent experiments. **E.** Comparison of the basal respiration, maximal respiration and the ATP-linked respiration of corrected and mutant hiPSC-CMs. N=10 wells for *LMNA* p.H222P and N=13 wells for Ctrl-iso 1, from 3 independent experiments. Mean ± SD. Mann-Whitney test. **F.** Measurements of ATP levels in LMNA and Crtl-iso hiPSC-CMs. Mann-Whitney test, *p<0.05. n = 8 from 3 independent experiments.

## DISCUSSION

Patients with cardiac laminopathy represent a group at high risk for developing heart failure and cardiac arrhythmias^2, 4^. However, the pathological mechanisms linking the nuclear envelope abnormalities induced by *LMNA* mutations to the development of a reduced cardiac muscle contractility are not well understood, limiting the development of targeted therapeutic approaches.

Here, we studied cardiomyocytes derived from hiPSCs generated from a patient carrying the p.H222P *LMNA* mutation^10^ and from a CRISPR/Cas9 genome-edited isogenic control. Cardiac differentiation of both cell lines resulted in cardiomyocytes with no or minor differences in their architecture and in the sarcomere assembly, but with significant differences in nuclear morphology which is a hallmark of *LMNA* mutations^5, 22^. Using a recently reported 3D cardiac organoid approach^14^, we were able to reproduce *in vitro* the reduced contractility found in patients with cardiac laminopathy, and these abnormalities were reversed by the genetic correction of the *LMNA* mutation. Interestingly, we found that the nuclear morphology was significantly changed in the 3D engineered cardiac tissues generated with the *LMNA* p.H222P mutant cardiomyocytes, and these changes were different from those observed in the 2D monolayer culture conditions. Nuclei in *LMNA* mutant 3D tissues were smaller and more elongated than their isogenic control counterparts, consistent with the *in vivo* observations in the cardiac tissue of *Lmna*^H222P/H222P^ transgenic mice^23^. Similarly, our work highlights the increased deformability of nuclei from *LMNA* mutant cardiomyocytes, suggesting an impaired mechano-transduction that is best revealed in 3D tissues that impose higher constraints from the surrounding cells^14^. To our knowledge, our study is one of the few to investigate 3D tissue engineering using hiPSC-CMs with *LMNA* mutations. Two studies showed that 3D engineered skeletal muscle using *LMNA*-mutant hiPSC-derived skeletal myotubes have an increased nucleus elongation^24, 25^. Moreover, a 3D environment was needed to reveal nuclear defects for the *LMNA* p.K32del and p.L35P mutations^24^.

Our work led us to discover a significant change in the MCUC stoechiometry and resulting abnormalities in mitochondrial calcium homeostasis in the *LMNA* mutant hiPSC CMs, which is one of the main findings of our study. These changes resulted from a significant reduction in the expression of MCU, the central pore on the inner mitochondrial membrane, and a significant increase in MICU1, a regulatory subunit that exerts an inhibitory effect at low calcium concentrations. The role of MCUC and mitochondrial calcium in the pathogenesis of heart failure remains controversial, as both up- and down-regulation have been shown to have deleterious effects on cardiac function. Indeed, *Mcu* deletion protected mice from cardiac dysfunction in the context of ischemia-reperfusion in two different studies^26, 27^, but others reported an improvement of cardiac function with a moderate overexpression of MCU in a model of heart failure in guinea pig^28^. In addition, kaempferol, an MCU activator, improved the arrhythmic phenotypes in mice with diabetic cardiomyopathy and hiPSC-CMs from cathecholergic ventricular tachycardic patients^29, 30^. It is likely that there is a biphasic effect of mitochondrial calcium on energetics, cardiac function and redox balance. Furthermore, the ratio between MCU and its regulatory subunit MICU1 has been found to control mitochondrial calcium uptake^20^. An increase in MICU1:MCU, as observed in our study, was found to decrease the mitochondrial calcium uptake and to support the development of cardiac dysfunction in mice^31^ and humans^30^. Consistent with these previous reports, we propose here a model in which changes in the MICU1:MCU leads to decreased mitochondrial calcium content, and reduced mitochondrial Ca^2+^ signaling that attenuates TCA cycle flux, compromising mitochondrial redox balance and ATP generation and leading to heart failure. The downregulation of MCU activity is consistent with our analyses of calcium transients which showed a paradoxical increase in the transient amplitude and area under the curve associated with a decreased calcium decay speed in the *LMNA* p.H222P mutant hiPSC-CMs^32, 33^. It also correlates well with the impaired mitochondrial respiration, and reduced ATP production. Moreover, increased MICU1:MCU associated with a decrease in ATP production would resolve the apparent contradiction between the increase in calcium transient amplitude and the decreased contractile forces.

*LMNA* mutations have previously been shown to have metabolic effects. Mutations leading to lipodystrophies or Hutchison-Gilford progeria syndrome (HGPS) induce metabolic disturbances, increased ROS and decreased mitochondrial respiration with an attenuated NAMPT-NAD pathway^34, 35^. Consistently, N-acetyl cysteine treatment or NR supplementation reduced cardiac oxidative stress injury and ameliorated contractile dysfunction in *Lmna*^H222P/H222P^ mice^8, 9^. Finally, impaired mitochondrial respiration has been reported in skeletal muscle cell lines carrying *LMNA* p.G232E or p.R482L mutations^36^. Here, we further report that the mitochondrial calcium uptake, which is critical for energy production and has been shown to play a role in the pathogenesis of cardiomyopathies^37^, is likely involved in the development of cardiac laminopathies. Our results indicate a significant effect of the *LMNA* p.H222P mutation on MCU mRNA levels, suggesting a change at the transcriptional level, as has been reported for other genes in the context of *LMNA*-related disorders. In addition, it remains to be tested whether the modulation of MCUC activity with pharmacological agents such as kaempferol or spermidine, or MCU overexpression *via* viral gene transfer could reverse the abnormal calcium transients and contractility observed in the *LMNA* mutant hiPSC-CMs. It was recently reported that the FDA-approved drug amorolfin could increase MCU-dependent mitochondrial metabolism in skeletal muscle and reverse muscle atrophy^38^. Whether this also applies to laminopathies remains to be demonstrated, but our study suggests that MCUC activators may represent a novel therapeutic option for patients presenting with *LMNA* cardiomyopathies.

Our study has some limitations. The 3D tissues were recorded under spontaneous pacing and unstressed conditions, which may lead to higher inter-tissue variability and explain the trend toward a bimodal distribution of contractility parameters observed in the *LMNA* p.H222P mutant cardiac tissues. Consistent with a mitochondrial dysfunction, it is likely that recording under pacing conditions would further accentuate the differences between groups, as observed with calcium transient recordings. Multiple mitochondrial functions are likely altered in the *LMNA* mutant hiPSC-CMs, as suggested by the deregulation of the expression of several genes involved in different mitochondrial processes. Additional experiments will be needed to further circumvent the mitochondrial abnormalities observed in patients with *LMNA*-cardiomyopathy.

In conclusion, we report the existence of a novel pathogenic mechanism linking mitochondrial dysfunction with impaired MCU complex activity, decreased mitochondrial calcium homeostasis, and reduced mitochondrial energy production to the reduced contractile dysfunction caused by the Lamin A/C mutation. Overall, our study opens new possibilities by suggesting MCU activators as a novel therapeutic for patients with *LMNA* cardiomyopathy.

## Acknowledgments

We thank the iPS core facility of Paris Cardiovascular Research Center for its support for cell culture. We thank Camille Knops and Yunling Xu from the Flow Cytometry and the Microscopy platforms, respectively, from Université de Paris, Paris Cardiovascular Research Center, Paris, France. Schemes were created with BioRender.com (access date:20/07/2024).

